## Supplementary Materials for "Unpacking the V1 map: Differential covariation of preferred spatial frequency and cortical magnification across spatial dimensions"

**Preferred spatial frequency (cycles/deg)**

| **Angle** | **Eccentricity** | | | | | | | |
|---|---|---|---|---|---|---|---|---|
| | **2°** | **3°** | **4°** | **5°** | **6°** | **7°** | **8°** | **9°** |
| **11.25°** | 2.00 | 1.70 | 1.41 | 1.21 | 1.10 | 0.98 | 0.87 | 0.77 |
| **33.75°** | 1.97 | 1.58 | 1.26 | 1.12 | 1.00 | 0.90 | 0.79 | 0.69 |
| **56.25°** | 1.89 | 1.53 | 1.18 | 1.01 | 0.92 | 0.83 | 0.76 | 0.68 |
| **78.75°** | 1.84 | 1.45 | 1.11 | 0.92 | 0.85 | 0.75 | 0.68 | 0.63 |
| **101.25°** | 1.97 | 1.61 | 1.19 | 1.00 | 0.86 | 0.78 | 0.72 | 0.68 |
| **123.75°** | 1.96 | 1.59 | 1.23 | 1.06 | 0.96 | 0.85 | 0.78 | 0.71 |
| **146.25°** | 2.00 | 1.63 | 1.32 | 1.12 | 1.01 | 0.90 | 0.82 | 0.74 |
| **168.75°** | 2.18 | 1.75 | 1.42 | 1.19 | 1.07 | 0.97 | 0.85 | 0.79 |
| **191.25°** | 2.30 | 1.83 | 1.48 | 1.25 | 1.11 | 0.98 | 0.90 | 0.80 |
| **213.75°** | 2.02 | 1.72 | 1.38 | 1.14 | 1.03 | 0.92 | 0.84 | 0.74 |
| **236.25°** | 1.90 | 1.55 | 1.26 | 1.08 | 1.00 | 0.91 | 0.83 | 0.73 |
| **258.75°** | 1.84 | 1.67 | 1.27 | 1.07 | 0.98 | 0.87 | 0.80 | 0.73 |
| **281.25°** | 1.93 | 1.56 | 1.24 | 1.06 | 0.94 | 0.85 | 0.82 | 0.78 |
| **303.75°** | 1.97 | 1.53 | 1.26 | 1.10 | 0.97 | 0.89 | 0.82 | 0.77 |
| **326.25°** | 2.01 | 1.61 | 1.35 | 1.15 | 1.05 | 0.96 | 0.88 | 0.79 |
| **348.75°** | 2.13 | 1.73 | 1.45 | 1.26 | 1.13 | 1.01 | 0.92 | 0.83 |

**Table S1: Group-average preferred spatial frequency (cycles/deg) for individual V1 segments.** Each polar angle bin is defined as 22.5° of angle centered at the labeled value (i.e., 11.25° of angle includes data between 0° and 22.5° of polar angle). 0° of angle is defined as at the right horizontal meridian of the visual field and angle increases counterclockwise around the visual field. Each eccentricity bin is defined as 1° of eccentricity centered at the labeled value (i.e., 2° eccentricity includes data between 1.5°–2.5° eccentricity). These values are derived from log-Gaussians fit to beta weights from the 'combined' stimuli condition (i.e. the average of the beta weights for each pinwheel-annulus stimulus pair.

**Cortical magnification (mm/deg)**

| Angle | Eccentricity | | | | | | | |
|---|---|---|---|---|---|---|---|---|
| | **2°** | **3°** | **4°** | **5°** | **6°** | **7°** | **8°** | **9°** |
| **11.25°** | 6.22 | 4.50 | 3.78 | 3.60 | 3.42 | 2.76 | 2.20 | 1.77 |
| **33.75°** | 5.71 | 4.31 | 3.37 | 2.96 | 2.62 | 2.20 | 1.83 | 1.48 |
| **56.25°** | 5.01 | 4.44 | 3.80 | 3.30 | 2.78 | 2.52 | 1.80 | 1.32 |
| **78.75°** | 3.96 | 3.07 | 2.95 | 2.33 | 1.94 | 1.79 | 1.49 | 1.07 |
| **101.25°** | 4.70 | 3.07 | 2.21 | 2.22 | 2.14 | 1.90 | 1.59 | 1.27 |
| **123.75°** | 5.68 | 3.96 | 3.38 | 3.03 | 2.59 | 2.21 | 1.90 | 1.36 |
| **146.25°** | 5.93 | 4.49 | 3.46 | 2.81 | 2.45 | 2.12 | 1.78 | 1.49 |
| **168.75°** | 6.56 | 4.80 | 3.83 | 3.13 | 2.85 | 2.49 | 2.22 | 1.89 |
| **191.25°** | 7.29 | 5.82 | 4.47 | 3.76 | 3.23 | 2.72 | 2.35 | 2.03 |
| **213.75°** | 6.63 | 4.97 | 4.20 | 3.31 | 2.66 | 2.42 | 2.00 | 1.48 |
| **236.25°** | 5.06 | 3.94 | 3.17 | 2.82 | 2.30 | 1.82 | 1.55 | 1.25 |
| **258.75°** | 5.00 | 3.56 | 2.65 | 2.62 | 2.48 | 2.07 | 1.82 | 1.44 |
| **281.25°** | 4.55 | 3.32 | 2.79 | 2.58 | 2.28 | 2.11 | 1.70 | 1.50 |
| **303.75°** | 5.01 | 3.44 | 3.21 | 2.97 | 2.47 | 2.02 | 1.79 | 1.47 |
| **326.25°** | 6.70 | 4.39 | 4.03 | 3.41 | 3.10 | 2.43 | 1.94 | 1.42 |
| **348.75°** | 6.70 | 5.25 | 4.72 | 3.70 | 3.17 | 2.89 | 2.34 | 1.98 |

**Table S2: Group-average V1 cortical magnification (mm/deg) for individual V1 segments.** Each polar angle bin is defined as 22.5° of angle centered at the labeled value (i.e., 11.25° of angle includes data between 0° and 22.5° of polar angle). 0° of angle is defined as at the right horizontal meridian of the visual field and angle increases counterclockwise around the visual field. Each eccentricity bin is defined as 1° of eccentricity centered at the labeled value (i.e. 2° eccentricity includes data between 1.5°–2.5° eccentricity).

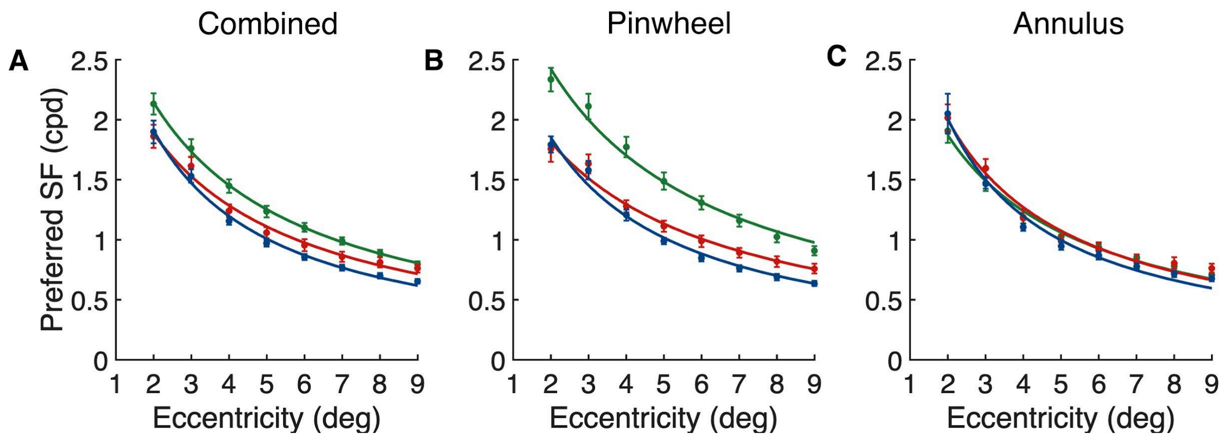


**Supplementary Figure S1: Polar angle asymmetries in preferred spatial frequency vary with stimulus orientation**. **(A)** Combined condition; preferred spatial frequency is highest along the horizontal, intermediate along the lower vertical, and lowest along the upper vertical meridian. **(B)** Pinwheel stimuli; the polar angle asymmetries are boosted as the pinwheel stimuli contain horizontal content along the horizontal meridian and vertical content along the vertical meridian. **(C)** Annulus stimuli; the polar angle asymmetries are weakened. The data are fit with an inverse linear function from [7]. Error bars represent ±1 standard deviation (SD) across 50 bootstrapped group-averages.

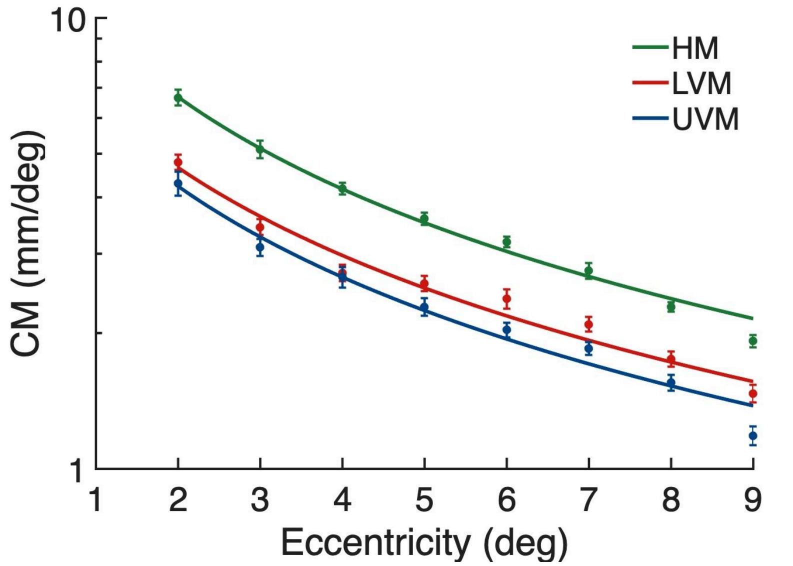


**Supplementary Figure S2. Polar angle asymmetries in V1 cortical magnification.** Cortical magnification plotted as a function of eccentricity for the horizontal meridian (HM: average of left and right horizontal), lower vertical (LVM), and upper vertical meridian (UVM). Data come from 22.5° wedge-ROIs centered either side of each meridian. The cortical magnification function from [7] is fit to the data from each meridian. Error bars represent ±1 SD across 50 bootstrapped group-averages.

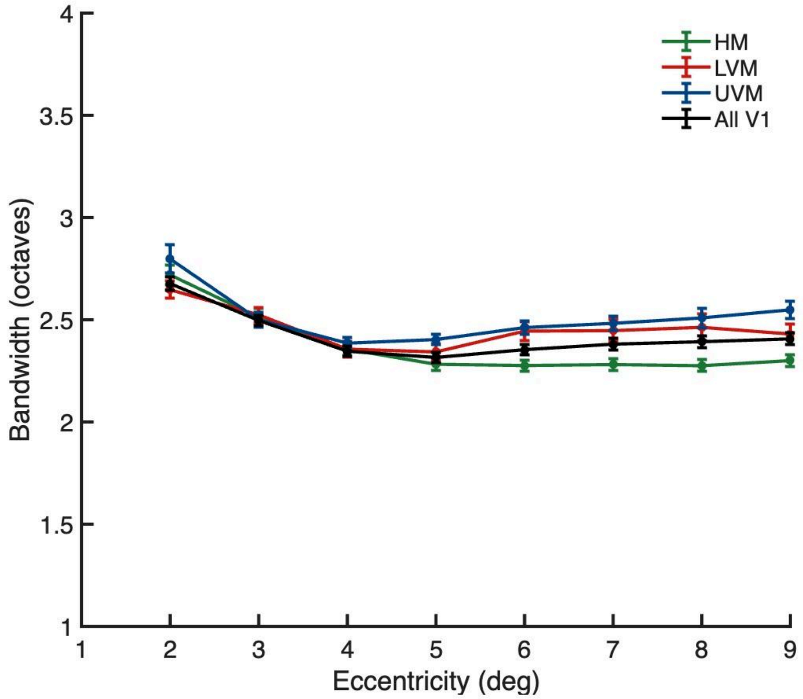


**Supplementary Figure S3: V1 spatial frequency bandwidth as a function of eccentricity.** Bandwidth (defined as σ of the log-Gaussian curve) varies as a function of eccentricity when measured along the horizontal, lower vertical, and upper vertical meridian, and all of V1 (combined stimulus condition). The meridian data are derived from 22.5° wedge-ROIs centered on either side of each meridian. All V1 data are averaged around polar angle. Bandwidth is indexed in octaves due to the logarithmic scaling of spatial frequency encoding in the visual system. Error bars represent ±1 standard deviation (SD) across 50 bootstrapped group-averages.

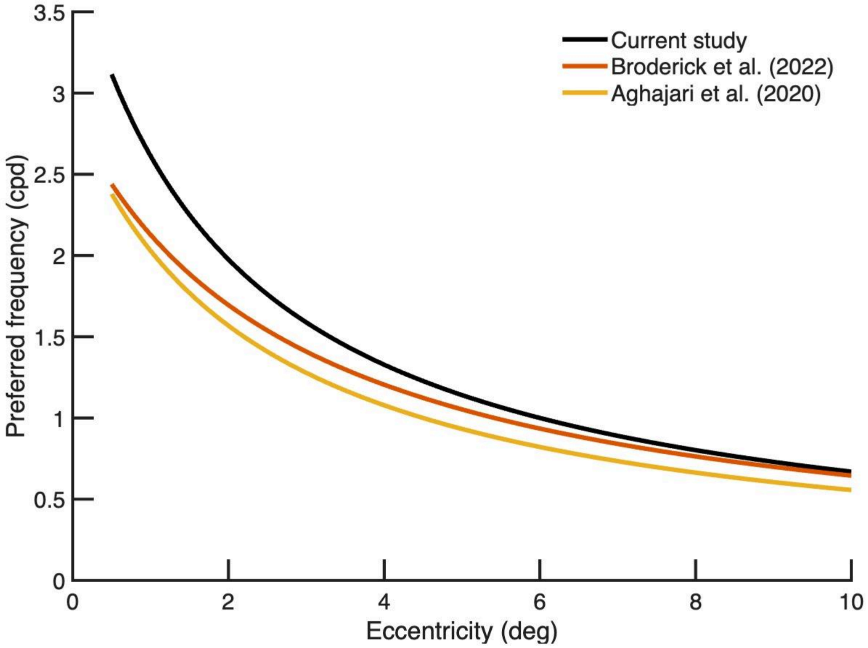


**Supplementary Figure S4. Comparing preferred spatial frequency as a function of eccentricity for scaled and uniform gratings.** Preferred spatial frequency is plotted as a function of eccentricity from two prior studies and current work. The current study and Broderick et. al. [11] fit the data with an inverse linear function, $F(R) = \frac{A}{(R+B)}$. We digitized the V1 data from Aghajari et. al. [12] Figure 4B and fit the same inverse linear function to their data. The three functions have slightly different shapes, however the estimates of preferred spatial frequency are close across the three studies.

**Median model parameter estimates when model is fit to individual observers**

| | Model parameter estimates | | | |
|---|---|---|---|---|
| | $A$ | $B$ | $\alpha$ | $\beta$ |
| **Preferred spatial frequency** | 8.6<br>$CI_{95}$ = [8.0, 9.3] | 2.4<br>$CI_{95}$ = [2.1, 2.7] | 0.09<br>$CI_{95}$ = [0.07, 0.11] | 0.04<br>$CI_{95}$ = [0.02, 0.06] |
| **Cortical magnification** | 17.3<br>$CI_{95}$ = [16.5, 18.1] | 1.3<br>$CI_{95}$ = [1.1, 1.6] | 0.20<br>$CI_{95}$ = [0.18, 0.25] | 0.03<br>$CI_{95}$ = [0.00, 0.06] |

**Supplementary Table S3.** Median model parameter estimates and 95% confidence intervals from the models fit to individual participant measurements of preferred spatial frequency and cortical magnification. 95% CIs are derived from bootstrapping across 40 observers.

*Supplementary analysis: Three anisotropies in spatial frequency tuning*

Here, we quantified how preferred spatial frequency varied between: 1) the horizontal and vertical meridian of the visual field; 2) radial and tangential stimulus orientations; and 3) horizontal and vertical stimulus orientations.

First, we examined how preferred spatial frequency varied between the horizontal and vertical meridian representation of the visual field, regardless of stimulus orientation. For each observer, we averaged together the preferred spatial frequency measurements for the pinwheels and annuli, with data localized with ±22.5° of either the horizontal or vertical meridian. Averaging across the stimulus classes for each location removes any asymmetry due to radial vs tangential or horizontal vs vertical. Preferred spatial frequency was 20–30% higher along the horizontal than vertical meridian of the visual field, after averaging out stimulus orientation (**FIG S5A**).

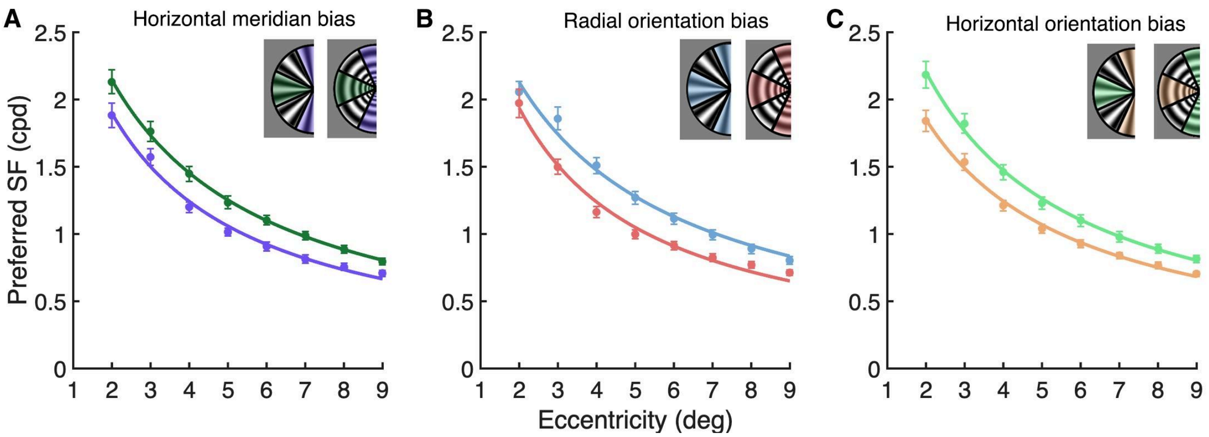


**FIG S5. V1 preferred spatial frequency varies with polar angle meridian and stimulus orientation**. **(A)** Preferred spatial frequency is higher along the horizontal than vertical meridian, after averaging out stimulus orientation. **(B)** Preferred spatial frequency is higher for radial (i.e., pinwheel stimuli) than tangential orientations (i.e., annulus stimuli), when averaged across the polar angle meridians. **(C)** Preferred spatial frequency is higher for horizontal than vertical stimulus orientations, after averaging out polar angle meridian. Error bars represent ±1 SD across 50 bootstrapped group-averages.

Second, we examined how preferred spatial frequency varied between radial and tangential stimulus orientations –irrespective of polar angle meridian. To this end, we computed preferred spatial frequency for the pinwheel (radial) and annulus (tangential) stimuli, averaged across location. For consistency, we restricted the locations to match those for the meridian analysis. Preferred spatial frequency was higher for radial than tangential stimuli, when averaged across the polar angle meridians, with the exception of the 2° eccentricity bin (**FIG S5B**). Note that averaging across the two meridians not only removes the horizontal vs vertical meridian asymmetry (**FIG S5A**), but also any horizontal vs vertical orientation asymmetry (**FIG S5C**).

Third, we tested how preferred spatial frequency varied between horizontal and vertical stimulus orientations. To compute preferred spatial frequency for horizontal stimulus orientations, we averaged data from the horizontal meridian for the pinwheels with data from the vertical meridian for the annuli, as both combinations have local horizontal orientation. For vertical stimulus orientations, we did the complement: we averaged data from the vertical meridian from the pinwheels with data from the horizontal meridian from the annuli. Preferred spatial frequency was systematically higher for horizontal than vertical stimulus orientations, after averaging out polar angle meridian (**FIG S5C**).

These anisotropies can interact. For example, the choice of stimulus would affect the result if one compared preferred spatial frequency across the different polar angle meridians. In **Supplementary FIG S2**, we present measurements of preferred spatial frequency as a function of eccentricity along the horizontal, lower vertical, and upper vertical meridian. When computed from the combined stimuli condition (**Supplementary FIG S2A**), preferred spatial frequency is highest along the horizontal, intermediate along the lower vertical, and lowest along the upper vertical meridian of the V1 representation. When computed from the pinwheel stimuli alone (**Supplementary FIG S2B**), the polar angle asymmetries increase. When computed from annuli alone (**Supplementary FIG S2C**), the polar angle asymmetries decrease.

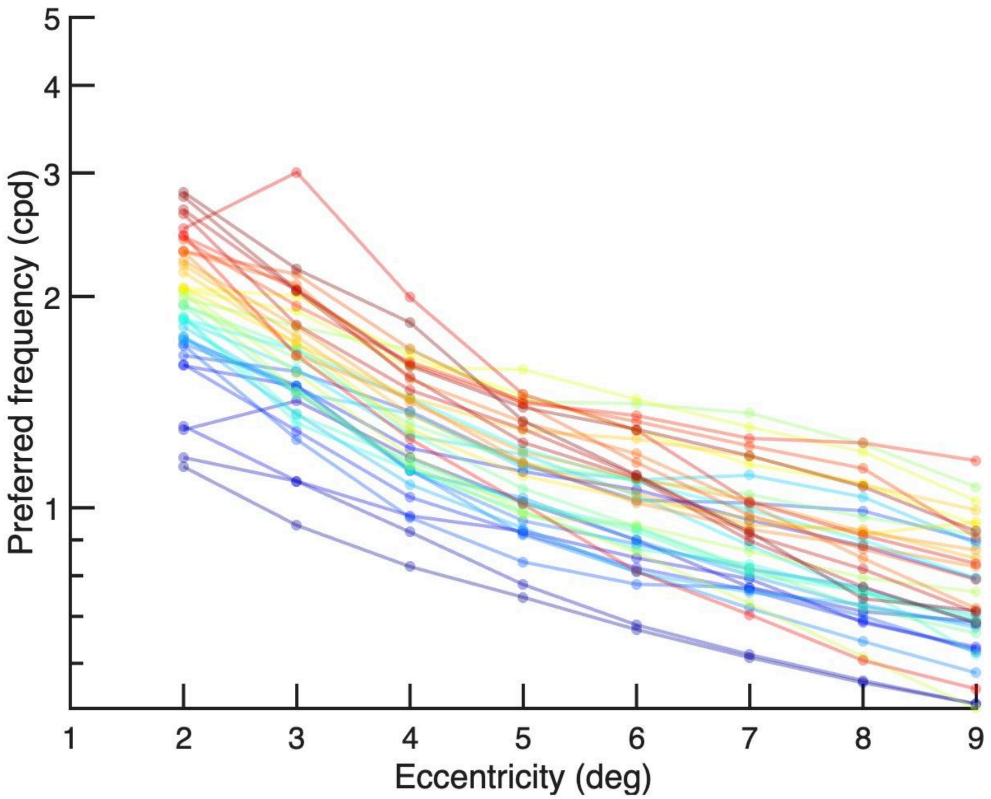


**Supplementary Figure S6. Preferred spatial frequency as a function of eccentricity for individual observers.** Each colored line represents the change in V1 preferred spatial frequency as a function of eccentricity for an individual observer (n=40).
